# RedRibbon: A new rank-rank hypergeometric overlap pipeline to compare gene and transcript expression signatures

**DOI:** 10.1101/2022.08.31.505818

**Authors:** Anthony Piron, Florian Szymczak, Maria Inês Alvelos, Matthieu Defrance, Tom Lenaerts, Décio L. Eizirik, Miriam Cnop

## Abstract

**Motivation:** High throughput omics technologies have generated a wealth of large protein, gene and transcript datasets that have exacerbated the need for new methods to analyse and compare big datasets. Rank-rank hypergeometric overlap is an important threshold-free method to combine and visualize two ranked lists of *P*-values or fold-changes, usually from differential gene expression analyses. Here, we introduce a new rank-rank hypergeometric overlap-based method aimed at both gene level and alternative splicing analyses at transcript or exon level, hitherto unreachable as transcript numbers are an order of magnitude larger than gene numbers.

**Results:** We tested the tool on synthetic and real datasets at gene and transcript levels to detect correlation and anti-correlation patterns and found it to be fast and accurate, even on very large datasets thanks to an evolutionary algorithm based minimal *P*-value search. The tool comes with a ready-to-use permutation scheme allowing the computation of adjusted *P*-values at low time cost. Additionally, the package is a drop-in replacement to previous packages as a compatibility mode is included, allowing to re-run older studies with close to no change to existing pipelines. RedRibbon holds the promise to accurately extricate detailed information from large analyses.

**Availability:** RNA-sequencing datasets are available through the Gene Expression Omnibus (GEO) portal with accession numbers GSE159984, GSE133218, GSE137136, GSE98485, GSE148058 and GSE108413. The C libraries and R package code are open to the community with a permissive licence (GPL3) and available for download from GitHub https://github.com/antpiron/ale, https://github.com/antpiron/cRedRibbon and https://github.com/antpiron/RedRibbon.

## 1 Introduction

During the past two decades, we have seen a democratization of high throughput sequencing technologies. The cost of DNA sequencing went down by six orders of magnitude from 2000 to 2022 (Lewin, et al., 2018). High throughput sequencing has led to the generation of large and diverse datasets covering multiple omics, including genomes, transcriptomes and proteomes. Alternative splicing generates massive protein diversity. Through the inclusion or exclusion of exons from pre-mRNAs, distinct mature mRNAs give birth to multiple proteins with different functions (Black, 2003). Thereby, in humans, more than 200,000 different proteins are produced from around 20,000 protein coding genes (Alvelos, et al., 2018). Alternative splicing is omnipresent in eukaryotic cells, with 80% of protein coding genes undergoing it. On average, a human gene is spliced into 4.4 transcripts. Alternative splicing is implicated in many diseases (López-Bigas, et al., 2005). Its analysis is challenging as most studies and pathway databases are gene-centric (Liberzon, et al., 2011) and the number of transcripts or splicing events can be overwhelming. Most studies aggregate transcript expression levels at gene level, thus losing crucial information about isoforms that may play different or even opposite roles. Collectively, the widespread use of these omics technologies has exacerbated the need for new methods to analyse and compare diverse and ever larger datasets. Multiple data aggregation initiatives collected those datasets, including amongst others: Gene Expression Omnibus (Barrett, et al., 2012; Edgar, et al., 2002), the Genotype-Tissue Expression (GTEx) Project (Lonsdale, et al., 2013) and the Translational Human Pancreatic Islet Genotype Tissue-Expression Resource (TIGER) (Alonso, et al., 2021).

Rank-rank hypergeometric overlap (RRHO) has been developed to compare two lists of differentially expressed genes generated with microarray technology (Plaisier, et al., 2010) and it was further improved with alternative statistics and better enrichment sets (Cahill, et al., 2018). RRHO compares two labelled ranked lists of real numbers. The labels can be gene/transcript identifiers or any other unique identifier. The values can be fold changes, *P*-values, slopes or another meaningful ranked statistic. The method detects the enrichments at the extremities of the ranked lists. For example, using two lists of fold change in gene expression, it allows the construction of enriched gene sets for the four possible directions, i.e. downregulated-downregulated, upregulated-upregulated, downregulated-upregulated and upregulated-downregulated. The method proceeds by computing, for all coordinates (*i, j*) in the two compared lists, an enrichment *P*-value from the number of labels in common at the extremities up to the coordinate with the hypergeometric distribution. The coordinates with the minimal *P*-value are used to determine the most significant gene set. The method can thus be seen as a 2D generalisation of Gene Set Enrichment Analysis (Mootha, et al., 2003; Subramanian, et al., 2005). The RRHO method allowed us and others to generate meaningful comparisons of differential gene and protein expression data (Blencowe, et al., 2022; Colli, et al., 2020; Colli, et al., 2020; Lytrivi, et al., 2020; Marselli, et al., 2020), but its application revealed shortcomings that required adaptation of the original RRHO R package (Marselli, et al., 2020). Nonetheless, major shortcomings remained. First, the original R package is limited by R language real number representation (R Core Team, 2022). This representation often leads to an underflow, i.e., a *P*-value that is rounded to zero below a threshold. Therefore, the zero *P*-values become indistinguishable from each other, making the detection of the minimal *P*-value impossible and lowering the accuracy of the method. Second, the execution time follows a cubic growth depending on the list length, making it unpractical for large lists. To circumvent these long run times, the original R package offers the possibility to skip some coordinates in the map, trading accuracy for performance. The recommended step size is between 100 and 500 for lists of 10,000 to 50,000 elements (with the number of element square root as default value) introducing a potential inaccuracy of hundreds of genes.

In recent years, methods have been developed to accurately quantify transcript expression levels, including splice variants, from RNA-sequencing (RNA-Seq) reads, e.g. Salmon (Patro, et al., 2017), kallisto (Bray, et al., 2016) and RSEM (Li and Dewey, 2011). Despite the progress in transcript quantification methods, to our knowledge, there are no tools available to compare transcript level differential analyses without prior gene level aggregation. As the total number of transcripts quantified by RNA-Seq is an order of magnitude higher than for genes, existing RRHO packages are inadequate. Plaisier et al. suggested to compute corrected *P*-values by permuting samples and re-running differential and RRHO analyses a thousand times (Plaisier, et al., 2010). This permutation method is slow for large gene expression analyses and renders transcript expression analyses prohibitive (or inaccurate with very large step sizes).

To address the above-described unmet need for comparative analysis tools of diverse omics datasets including transcripts, we developed RedRibbon. RedRibbon is a complete rewrite of the original RRHO package bearing in mind performance and accuracy, introducing novel data structures and algorithms, and an all-in-one permutation method to adjust the minimal *P*value. The improvements in performance and accuracy have been assessed using synthetic datasets and previously reported results (Marselli, et al., 2020). We applied the method to compare alternative splicing results in experimental models of diabetes, including the human EndoC-βH1 beta cell line and pancreatic islets.

## 2 Methods

The RedRibbon package is a complete rewrite of the original package (Plaisier, et al., 2010; Rosenblatt and Stein, 2014) with performance in mind and including novel algorithms and data structures. RedRibbon allows to do a full RRHO analysis with adjusted *P*-values over lists or differential analyses containing millions of elements. A *C* library and an easy-to-use *R* package are provided. It implements new performant algorithms to find the minimal *P*-value, a novel adjusted minimal *P*-values computation algorithm, improved plots and parallel execution.

### 2.1 RedRibbon workflow

The input for RedRibbon is two lists of gene or transcript ranked statistics such as fold change or direction signed P-value (Fig. 1A). From these, the minimal hypergeometric P-value coordinates are identified in the four quadrants of the level map using the original method (Plaisier, et al., 2010) – called in this manuscript “grid method” in reference to the grid-like traversal of the coordinate matrix – or our fast and accurate evolutionary algorithm (see below). Locating the minimal coordinates aims to split overlapping map quadrants into two areas, an enriched and a randomly ordered region. Optionally, the minimal P-value can be adjusted considering expression level correlation between genes or transcripts. The result is four transcript sets, one per quadrant. The enrichment result, the corrected P-values and the four quadrants can be visualized in an overlap map. Pathway enrichment analyses usually use overlapping genes in each quadrant. For transcript level analyses, as most databases are gene centric, it is necessary to convert transcripts to genes. Gprofiler2 R package (Kolberg, et al., 2020) and clusterprofiler R package (Wu, et al., 2021; Yu, et al., 2012) were used to do the enrichment analysis respectively for gene level and alternative splicing analyses. For alternative splicing analysis, we created 5 new pathways, namely SRSF6 regulation, type 2 diabetes, positive regulation of apoptosis, insulin secretion and JNK signalling. The list of spliced genes regulated by SRSF6 were taken from (Juan-Mateu, et al., 2018) (the spliced gene lists are reproduced in the present Fig. 4A) and have been converted into clusterProfiler-ready format.

**Fig. 1.**
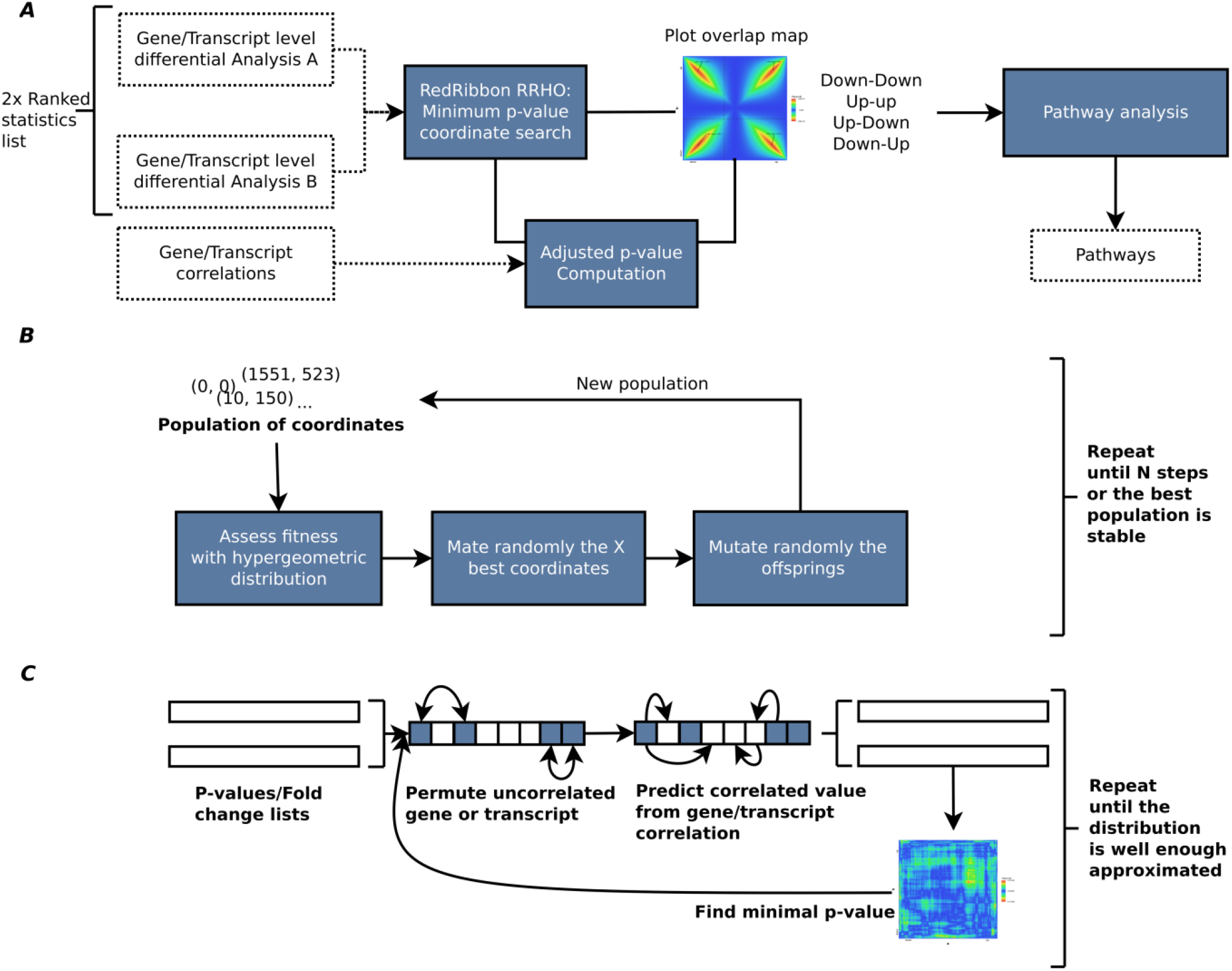
RedRibbon RRHO workflow. (A) Transcript level differential analysis by RRHO. The RedRibbon RRHO package can handle very large data because of improved data structures and algorithms (see benchmark). Transcript level differential analysis can be overlapped with a permutation scheme to correct *P*-values. The overlap analysis is followed by a pathway analysis. (B) The evolutionary algorithm will find the minimal *P*-value among coordinates. The best fitness individuals of a population of coordinates are mated and then randomly mutated to obtain a new population. This process is repeated until stability is reached among the best population or a fixed number of steps. (C) Hybrid prediction-permutation method to compute the adjusted minimal *P*-value. A set of uncorrelated elements (genes, transcripts; shown in blue squares) is identified. Their value (*P*-value or fold change) is permuted. The remaining correlated elements of the lists are predicted from this set with a linear model. The minimal RRHO *P*-value is then computed for the two permutated lists. The operation is repeated a fixed number of times and the adjusted *P*-value assessed.

### 2.2 RedRibbon rank-rank hypergeometric overlap

#### 2.2.1 P-value computation

RedRibbon *P*-values can be computed with oneor two-sided or compatible with the original R module two-sided statistics. With *c* being the number of genes in common for the coordinate (*i, j*) in an RRHO map of size *n* × *n*, the *P-value* is computed as

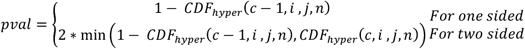

The two-sided test allows to detect enrichment both in correlated (up/up and down/down) and anti-correlated (up/down and down/up) genes. The anti-correlated genes were not reported by the original R Package. Additionally, if the hypergeometric *P*-value comes from the lower tail of the distribution, they are negatively signed to distinguish depletion (anti-correlation) from enrichment (correlation).

#### 2.2.2 C language implementation and R module

RedRibbon has been split between a performant C library and an easy-touse R module interfacing this library. RedRibbon has been optimised to be efficient regardless of the length of the lists given as input. The gene sets are represented by bit vectors allowing the use of CPU bit instructions. Bit vectors allow to efficiently compute set intersections, an essential operation for RRHO as it is done for each *P-value* computation. Additionally, the intersected gene sets relative for two close coordinates on the RRHO map are very similar, most genes being in common. We leverage this similarity to decrease the operation numbers to compute the intersections by updating previously computed sets.

In order to improve the accuracy of small *P-value* computation which can be smaller than the smallest representable positive number by C language, *long double* type has been used for large lists rather than *double*. Doing so, on a x86 platform, the smallest positive number becomes 3.36*e*^−4932^ instead of 2.23*e*^−308^. As *R* does not support the *long double* type, we use the logarithm of the *P-value* to stay in the representable range for the *R* package.

### 2.3 Evolutionary algorithm to find the minimal P-value

Two minimal *P-value* search methods are implemented in RedRibbon: (1) the grid-based method used in the original RRHO implementation, and (2) an evolutionary algorithm-based method (Fig. 1B). The latter interprets coordinates on the map as individuals of a population subject to selective pressure. The initial set of coordinates is chosen to be uniformly spaced on the diagonal of the RRHO map, a choice purposely reminiscent of the classical method to guarantee with a high probability that it remains at least as good as the classical method. Next, a new generation is created by mating the coordinates and randomly introducing mutations to induce genetic diversity. This new population is then selected for the coordinates with the best fitness, measured with hypergeometric *P*-values and where lower is better. The process is repeated until a stable set of best coordinates (i.e., no newly added coordinates in best coordinate set) or a pre-defined number of generations is reached.

### 2.4 Adjusted minimal P-values

The RRHO minimal *P-value* coordinate is selected among a large set of coordinates. For lists of N features, the minimal *P*-value coordinate is to be found in N^2^ coordinates, each associated with one colour dot in the overlap map. Correcting the minimal coordinate *P*-values presents multiple challenges: (1) while the distribution for one specific coordinate is hypergeometric, the distribution of the minimal *P-value* coordinate is, to our knowledge, unknown, (2) the features can be correlated, e.g. from gene interaction, (3) the coordinate *P*-values are highly dependent as the hypergeometric *P-value* for two close coordinates is computed from sets with many elements in common; hence, the false discovery rate correction assuming independence of variables is inadequate to compute a corrected minimal *P-value*, (4) permuting samples and re-doing the full differential expression analyses many times as in Plaisier et al. (Plaisier, et al., 2010) is very time consuming, unpractical and out of reach for very large datasets. Here, we introduce a new permutation scheme considering the correlation without re-doing the differential analysis for each permutation (Fig. 1C).

#### 2.4.1 Hybrid prediction-permutation method

RedRibbon adjusted minimal *P*-values are computed using an efficient hybrid prediction-permutation method (HPP) to assess the null distribution of the minimal coordinate *P-value*. When the features are independent, the HPP method is strictly equivalent to a permutation of the feature lists. The HPP method divides the features in two disjoint sets, namely the predictor and predicted sets. The predictor set is composed of features which are not able to mutually predict each other. Each predicted set feature value is predictable from the predictor set feature values or another predicted feature. A HPP permutation is generated by first permuting the original predictor set values and then predicting the other set features from these.

For a list of fold changes *FC*_*i*_, the predicted values are computed with a linear model:

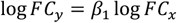

where the gene or transcript *x* is in the predictor set and *y* belongs to the predicted set and the *β*_1_ coefficient is estimated from another linear model over expression levels log *Expr*_*y*_ = *β*_1_ log *Expr*_*x*_ + *β*_0_. The latter is justified as for two samples we have 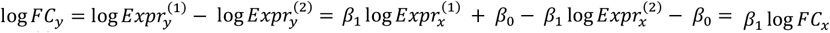. The linear model parameters are estimated with ordinary least square method from the expression matrices.

For a list of *P*-values *P*_*i*_, the predicted values are computed from the expression correlation coefficient *r* relative to a predictor and a random value 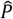 generated from the distribution of all values in the original list:

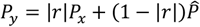

where *P*_*x*_ is in the predictor set and *P*_*y*_ belongs to the predicted set.

This formula assumes a linear effect between the value and the correlation coefficient. This effect is modelled by the term |*r*|*P*_*x*_. In case of standardised predictor and predicted variables r is the best linear ordinary least square estimator. The term 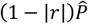 is a bootstrap estimate before taking into account the correlation of variables. The whole formula guarantees that the estimated value is equal to the predictor value if |r| is one while an r close to zero gives a bootstrapped random value. Consequently, the model can be seen as a finite mixture between the predictor and a bootstrap variable weighted with the correlation coefficient.

#### 2.4.2 Beta distribution fitting

The HPP method is repeated several times to obtain a list of minimal *P*-values. In order to limit the number of HPP iterations (around 100), a beta distribution is fitted on the HPP *P*values, and the goodness of fit is assessed with a Kolmogorov–Smirnov test. If the goodness of fit test is verified, the threshold for 0.05 significance is given by moment methods fitted beta *CDF*^−1^(0.05) and the adjusted *P*-value is 0.05/(*CDF*^(−1) (0.05)) ∗ *pvalue*. If the goodness of fit test rejects the hypothesis of beta distribution, the threshold is computed from the empirical cumulative distribution function.

#### 2.4.3 Computing the parameters of linear regressions and selecting HPP predictors

In case of fold change lists, for each gene the best gene predictor is selected among the linear regression models with significant *β*_1_. The minimal mean squared error model is used as fold change predictor. A gene is put in the HPP predictor set if it is the best predictor for another gene and is not itself predicted from another gene. For one gene, all linear models are computed at once with an efficient ordinary least square based on Householder reflections QR decomposition. For *P*-value lists, the best significant |*r*| is selected.

### 2.5 Data structures and algorithm

The computation of hypergeometric enrichment requires the computation of many set intersections. A bitset data structure is used to represent the sets. The bitset is a C array of 64 bits unsigned integer allowing to intersect 64 elements in one computer cycle with a binary “*and”* instruction. This data structure divides the number of operations for the intersection by 64 and lowers the CPU cache pressure (Drepper, 2007) reducing the RAM storage by 32, the integer bit size of R language.

Additionally, the intersection algorithm makes use of the previous intersection computation to reduce the number of updates to sets to intersect as the sets for the coordinate (*i* + *b, j*) are the same as for previously computed (*i, j*) set except for *b* elements.

### 2.6 Synthetic gene sets and accuracy measurements

In order to assess the accuracy of our method, two distinct synthetic sets of gene sets have been generated. The aim of these datasets is to assess the true positive rate (i.e., sensitivity), true negative rate (i.e., specificity) and the accuracy in a setting where the overlapping genes are known. The first set of gene sets (TS1) is composed of 192 artificial gene sets – each of 23,000 genes with 5000 genes with perfectly identical fold changes in both lists, split in half between downregulation and upregulation. The remaining genes have randomly assigned fold changes. This set has no noise for the 5000 overlapping genes. Hence, the boundaries between down- and upregulated genes are well defined, and the overlapping methods are expected to detect close to perfectly the 5000 overlapping genes.

The second set of gene sets (TS2) is composed of 192 artificial gene sets of 23,000 genes with 5000 log fold change correlated genes like l*FC*_*b*_ = *sign*(l*FC*_*a*_) ∗ *X*(|l*FC*_*a*_|^−1^) for each gene, where *X*(*λ*) is an exponential random variate of *λ* mean, *a* and *b* are the two lists. The *lFC*_*a*_ and uncorrelated *lFC*_*b*_ are generated from a standard normal distribution. The noisy association between the 5000 genes is closer to real data. Hence, the boundaries between down- and upregulated genes are not as well defined and are harder to detect with a rank-rank hypergeometric overlap.

### 2.7 Transcriptomes and differential analyses

RNA-Seq datasets previously generated by our group (see structured abstract, availability section) were used to assess the performance of RedRibbon, and its ability to generate new and accurate results. The data are transcriptomes of human islets of Langerhans from type 2 diabetic and non-diabetic donors, the latter exposed to palmitate and high glucose for 2 days followed by a recovery period from the lipoglucotoxic insult of 4 days (Marselli, et al., 2020), or to IFNα for 8 and 18h (n = 6) (Colli, et al., 2020; Gonzalez-Duque, et al., 2018), and EndoC-βH1 cells – an immortalized human beta cell line – exposed to IFNα for 8 and 18h (n = 5) (Colli, et al., 2020), IFNγ+IL-1β for 24h (n = 5) (Ramos-Rodriguez, et al., 2019) or following knockdown of the splicing factor *SRSF6* (n = 5) (Juan-Mateu, et al., 2018).

Quality control and trimming were done with fastp 0.19.6. The bulk RNASeq fastq were quantified with Salmon 1.4.0 (Patro, et al., 2017) using the parameters *--seqBias --gcBias –validateMappings* with GENCODE v36 (Frankish, et al., 2019) as the genome reference. Differential expression analyses were done with DESeq2 1.28.1 (Love, et al., 2014).

### 2.8 RedRibbon R package compatibility with the original implementation

In order to facilitate re-analysis of existing datasets, the RedRibbon R package provides a compatibility mode. The original *RRHO* R function has been rewritten using the new algorithms of RedRibbon. Hence, the existing pipelines can be improved for accuracy and performance just by substituting the library inclusion with close to no code editing.

## 3 Results

### 3.1 Enhanced overlap maps

We first generated synthetic dataset overlap maps to exemplify RedRibbon results and illustrate its new visual features that facilitate interpretation (Fig. 2). The overlap map of two perfectly identical lists is a perfect diagonal signal from downregulation to upregulation (Fig. 2A). The hypergeometric *P-value* gets lower as coordinates are closer to the list centres and give the whole list as enrichment. On the contrary, the overlap map of one list being in perfectly reversed order of the other — i.e., perfect anti-correlation of gene expression changes — follows a perfect diagonal from down-up quadrant to the up-down quadrant (Fig. 2B). The *P*-values are negatively signed in order to distinguish them as related to anti-correlated genes. The P-value can be plotted with different colours in the overlap map depending on their sign (not shown here).

**Fig. 2.**
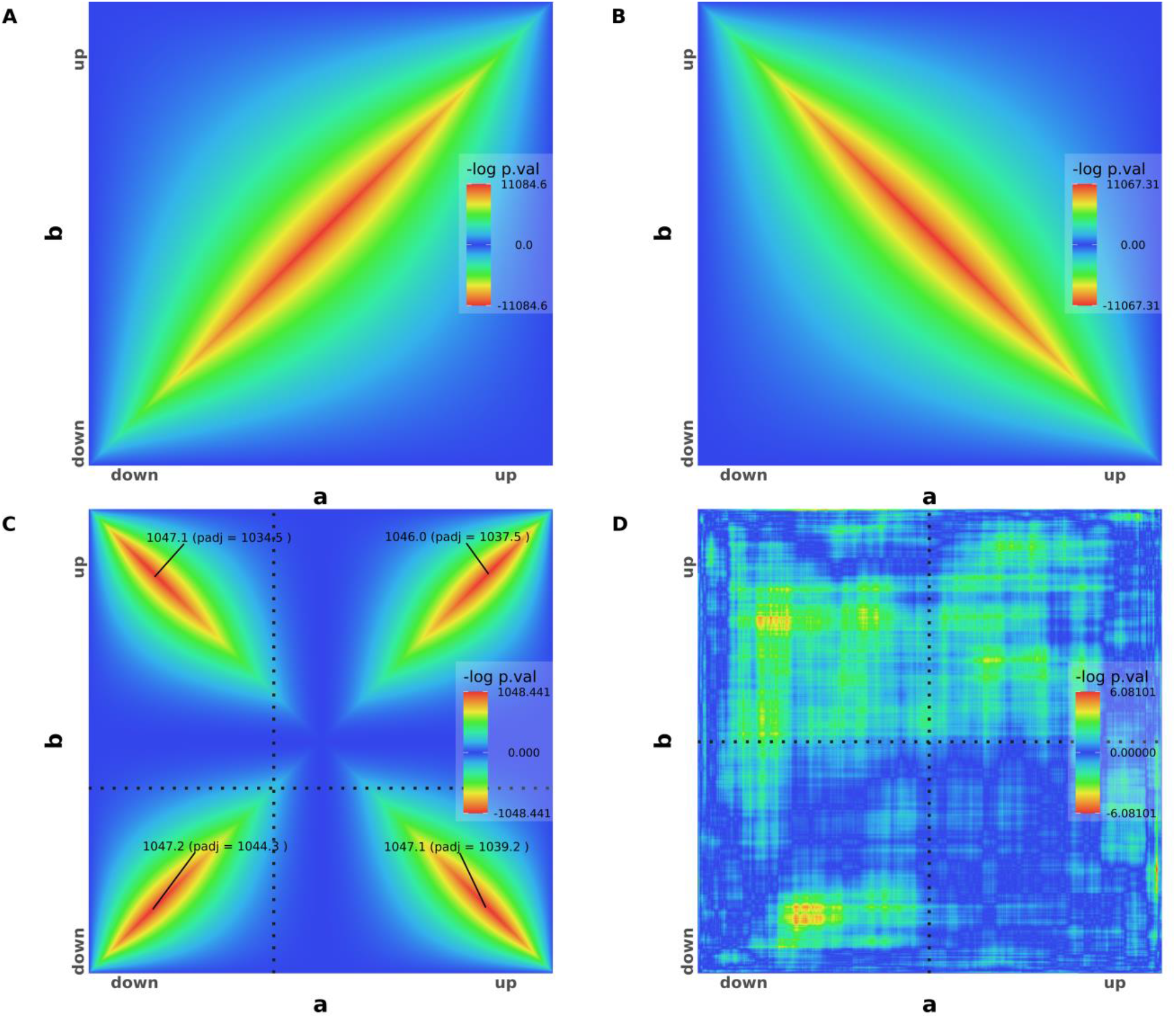
Rank-Rank Hypergeometric Overlap for artificial datasets. (A) RRHO map of two perfectly identical lists a (on the x-axis) and b (on the yaxis). (B) RRHO map of two perfectly symmetrical list going in opposite direction. (C) Two lists with half of the elements going in the same direction and the other half in opposite direction. The 96 permutation-adjusted *P*-values are reported. (D) RRHO map of two random lists. All adjusted *P*-values are below the significance threshold and therefore not shown (greater than 3 ≈ − log(0.05)).

Two lists with four quarters of 5000 genes going respectively and perfectly in the same direction (both down-ranked or both up-ranked in the lists) or in opposite directions (down-ranked in the first list and up-ranked in the other one, and vice-versa) result in an overlap map with perfect diagonal signals for the four quadrants (Fig. 2C). The maximal log *P*-values and the permutation adjusted *P*-values are shown for each quadrant. The horizontal and vertical dotted lines split the downregulation and upregulation where the log fold change is zero. In this dataset, the “zero” log fold change is at two fifth of both lists, hence, the split point is shifted to the beginning of the lists.

For completely random synthetic data lists, the fluctuation in the map is caused by random sampling and no signal is present (Fig. 2D). In this case, the overlap algorithm is unable to find any significant adjusted *P*-value and no *P*-value is shown on the map for any quadrant.

### 3.2 Performance

We next benchmarked RedRibbon against the original R package (Fig. 3). First, both packages were compared using the same grid method for a list of n genes with a step size of 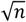, aiming to assess the performance of the new data structures and intersection algorithm (see Methods). On an Intel® Xeon® Processor E5-2650 v4, RedRibbon’s running time increases slowly with the gene list size and is below 25 seconds for lists of 262,144 genes, while the running time of the original R implementation grows steeply and is already close to 200 seconds for lists of 65,536 genes (Fig. 3A, left).

**Fig. 3.**
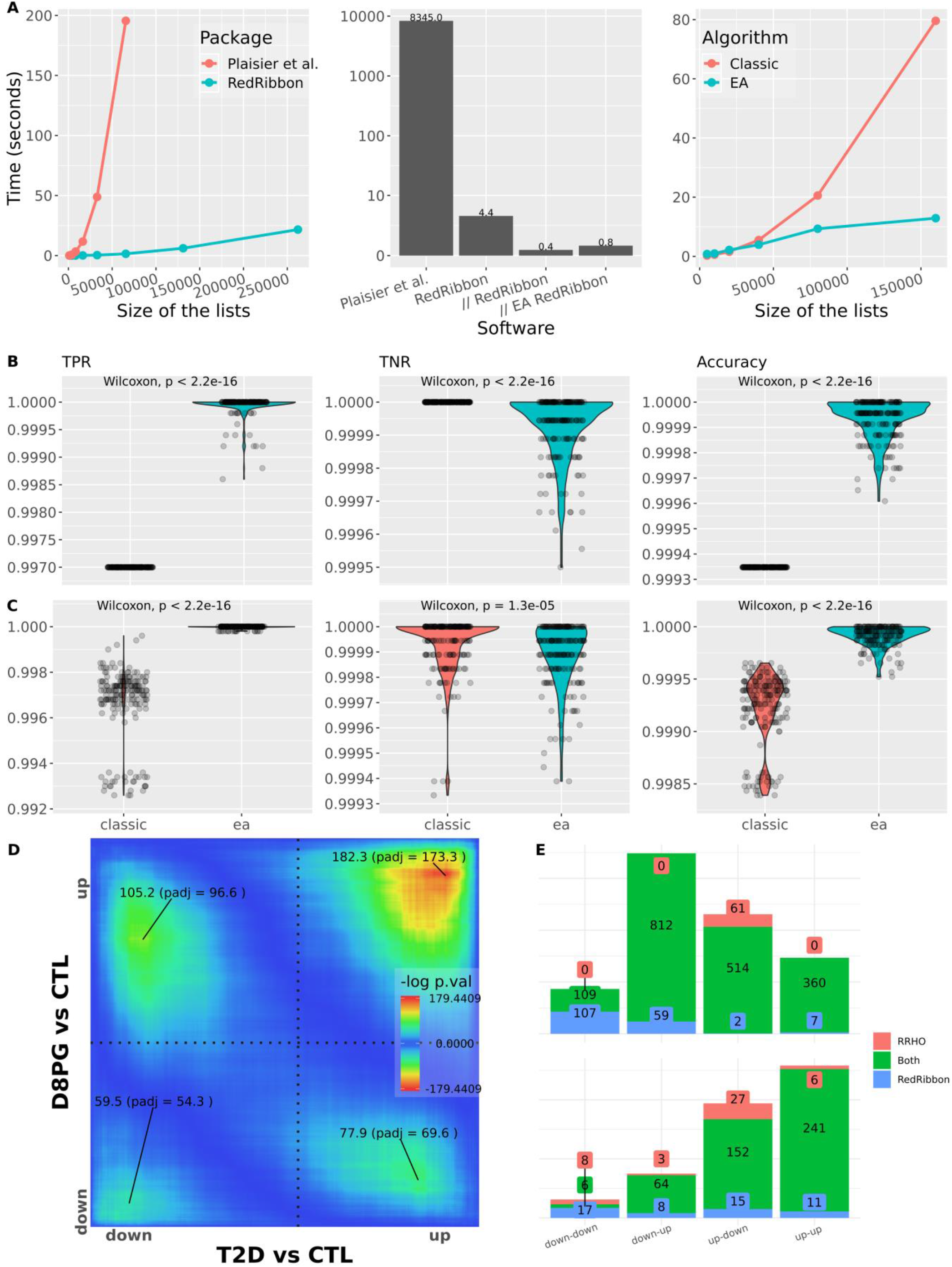
Benchmark of RedRibbon. (A) Assessment of RedRibbon performance. Left: Time to compute the minimal *P*-value for n genes with a step size of sqrt(n) with the Plaisier et al. (Plaisier, et al., 2010) grid method compared to our re-implementation of this method using the same parameters. Center: Comparison of our *P*-value permutation method with Plaisier et al. for 5,000 genes and a step size of 50. Time according to Plaisier et al. is reported and corrected for CPU performance improvement (single thread performance on https://www.cpubenchmark.net/). The RedRibbon method is reported in single thread (RedRibbon), multithreads (// RedRibbon), and multithreads with the evolutionary algorithm (// EA RedRibbon). Right: Time to compute the minimal *P*-value of n genes with a step size of sqrt(n) with permutation *P*-value correction. Our re-implementation of the Plaisier et al. grid algorithm is compared to the new evolutionary algorithm that has no step size limitation and hence higher accuracy. (B) Results for 192 artificial datasets of 23,000 genes with 5000 genes with perfectly identical fold changes in both lists, split in half between lowest and highest fold change (see method test set TS1). The remaining genes have randomly assigned fold changes. True Positive Rate (TPR), True Negative Rate (TNR) and accuracy are reported. TPR and accuracy are significantly better for evolutionary algorithms (ea) than with the classic grid method. (C) Violin plots for 192 artificial datasets of 23,000 genes with 5000 log fold change correlated genes (see method test set TS2). (D) RedRibbon hypergeometric map of >16,547 genes comparing human islets from type 2 diabetic (T2D) vs non-diabetic (CTL) donors and human islets recovering from palmitate+glucose exposure in vitro (D8PG vs CTL), as in (Marselli, et al., 2020). (E) Comparison of the original RRHO algorithm with default step size (128 genes, red) with RedRibbon (blue). Green shows elements detected with both methods. Top figure shows the overlapping gene counts. Bottom figure shows pathway enrichment counts.

Next, two lists of 5,000 genes and a step size of 50 were used to assess the *P*-value adjusting method, a benchmark setting used by Plaisier et al. (Plaisier, et al., 2010). The running time they reported (8,345 seconds, after correction for CPU performance) is used as reference. RedRibbon outperforms this by four orders of magnitude for all tested methods: grid method, parallel execution grid, and evolutionary algorithm (Fig. 3A, middle).

Our grid method re-implementation was then compared to the evolutionary algorithm with adjusted *P*-value computation. For the grid method, we used a square root of the list length for the step size. Both algorithms were run in parallel mode for the adjusted *P*-value permutation computation. The grid method outperforms the evolutionary algorithm for shorter lists, but with 20,000 or more genes the evolutionary algorithm has an advantage (Fig. 3A, right). The evolutionary algorithm is usable for the analysis of millions of elements (up to 256,000 are shown in Fig. 3A).

### 3.3 Accuracy

True positive rate (TPR), true negative rate (TNR) and accuracy were assessed for the two synthetic datasets (see Methods). Measurements for the first set show a clear-cut advantage to the evolutionary algorithm for both TPR and accuracy (Fig. 3B). In most synthetic datasets, TPR is exactly 1 meaning that all genes significantly correlated in the lists are detected, while for the grid method 3 genes in 1,000 are missed (TPR = 0.997). The TNR is kept under control as at most 0.5 gene out of 1,000 misses detection (TNR = 0.9995) with most datasets having close to 0 misdetections. Accuracy is systematically better for the evolutionary algorithm than for the grid algorithm (Fig. 3B).

Measurements for the second synthetic dataset — simulating a real case scenario — is clearly to the advantage of the evolutionary algorithm for TPR and accuracy (Fig. 3C). TPR is equal or close to 1 for all generated gene sets. The grid method TPR and accuracy exhibit a bimodal distribution caused by the step size length jumps in the coordinate. The TNR is close to a value of 1 as for the first dataset.

We next assessed RedRibbon on experimental datasets previously analysed by us (Marselli, et al., 2020). Rank-rank hypergeometric overlap was run between the fold changes of 16,547 genes from human islets, comparing donors with and without type 2 diabetes against islets exposed in vitro to the saturated free fatty acid palmitate and high glucose for 48h and subsequently allowed to recover for 4 days. The level map shows significant signals in the four quadrants with the strongest signal being in the upregulated direction (Fig. 3D). The comparison with the original R package shows a large intersection between the result of the original R and RedRibbon packages (Fig. 3E). RedRibbon identifies 7 to 107 additional genes in the 4 quadrants of the overlap map (Fig. 3E, top). The differences between the 2 packages result in differences in enriched pathway detection (Fig. 3E, bottom). The extent of the differences is similar to the differences for the synthetic gene sets, suggesting the accuracy metrics are sound.

### 3.4 Adjusted *P*-Value type 1 error

The permutation method was controlled for type 1 error against a random background composed of 1000 random list pairs of 1000 elements. An adjusted P-Value below 0.05 was reported for 1.3 percent of the RedRibbon analyses (*P*-Value = 2.5e-10 for *P*-Adjusted > 0.05 null hypothesis). This below expected percentage shows that our adjustment method conservatively controls for type 1 errors.

### 3.5 Alternative splicing analyses

The tool developed by Plaisier et al. is not suited for splicing analyses, while RedRibbon allows it by having the power to run hundreds of thousands of transcripts (Figure 3A). To validate the suitability of this package for this type of analysis, we applied RedRibbon to previously generated alternative splicing data. We have used our previously published RNASeq from EndoC-βH1 cells and human islets exposed to IFNα, and RNASeq in which the splicing factor *SRSF6* (also known as *SRp55*) was silenced (Juan-Mateu, et al., 2018). The exposure to IFNα induces beta cell hallmarks of type 1 diabetes, including inflammation, endoplasmic reticulum stress and HLA class I overexpression (Coomans de Brachène, et al., 2018; Marroqui, et al., 2017), but also major alterations in the splicing pattern (Colli, et al., 2020).

Interestingly, the main *SRSF6* transcript is downregulated in EndoC-βH1 cells and human islets exposed to IFNα (see Supplemental Table 1, transcript *SRSF6-201*). *SRSF6* seems to be thus responsible for some of the IFNα-induced alternative splicing modifications (Juan-Mateu, et al., 2018), providing an interesting model for comparison with splicing in *SRSF6*-silenced EndoC-βH1 cells. *SRSF6* downregulation modulates the splicing of genes involved in apoptosis, JNK signalling, insulin secretion and type 2 diabetes (Fig. 4A from the results of (Juan-Mateu, et al., 2018)). The transcript signatures of *SRSF6*-depleted (101,226 transcripts) vs IFNα-exposed EndoC-βH1 cells (151,157 transcripts) show substantial overlap in down- and upregulated transcripts (Fig. 4B, left panel). The enrichment of SRSF6-regulated pathways and the type 2 diabetes pathway observed among the downregulated transcripts, points to a SRSF6mediated splicing modification in IFNα-treated EndoC-βH1 cells (Fig. 4C, left panel). The down-up and up-down overlap corresponds to changes induced by IFNα that are not recapitulated by SRFS6 knockdown and vice-versa, not discussed as we focus here on similarities. The comparison between EndoC-βH1 cells and human islets (165,066 transcripts), both exposed to IFNα, shows strong similarity in down- and upregulated transcripts, with an overlap pattern resembling Figure 2A, suggesting that EndoC-βH1 cells are an adequate model for alternative splicing studies (Fig. 4B, right panel). Among the downregulated transcripts, pathway analysis identified enrichment of the SRSF6 regulatory network – apoptosis, JNK signalling, insulin secretion, and type 2 diabetes – suggesting that SRSF6regulated splicing modifications are also present in human islets exposed to IFNα (Fig. 4C, right panel).

**Fig. 4.**
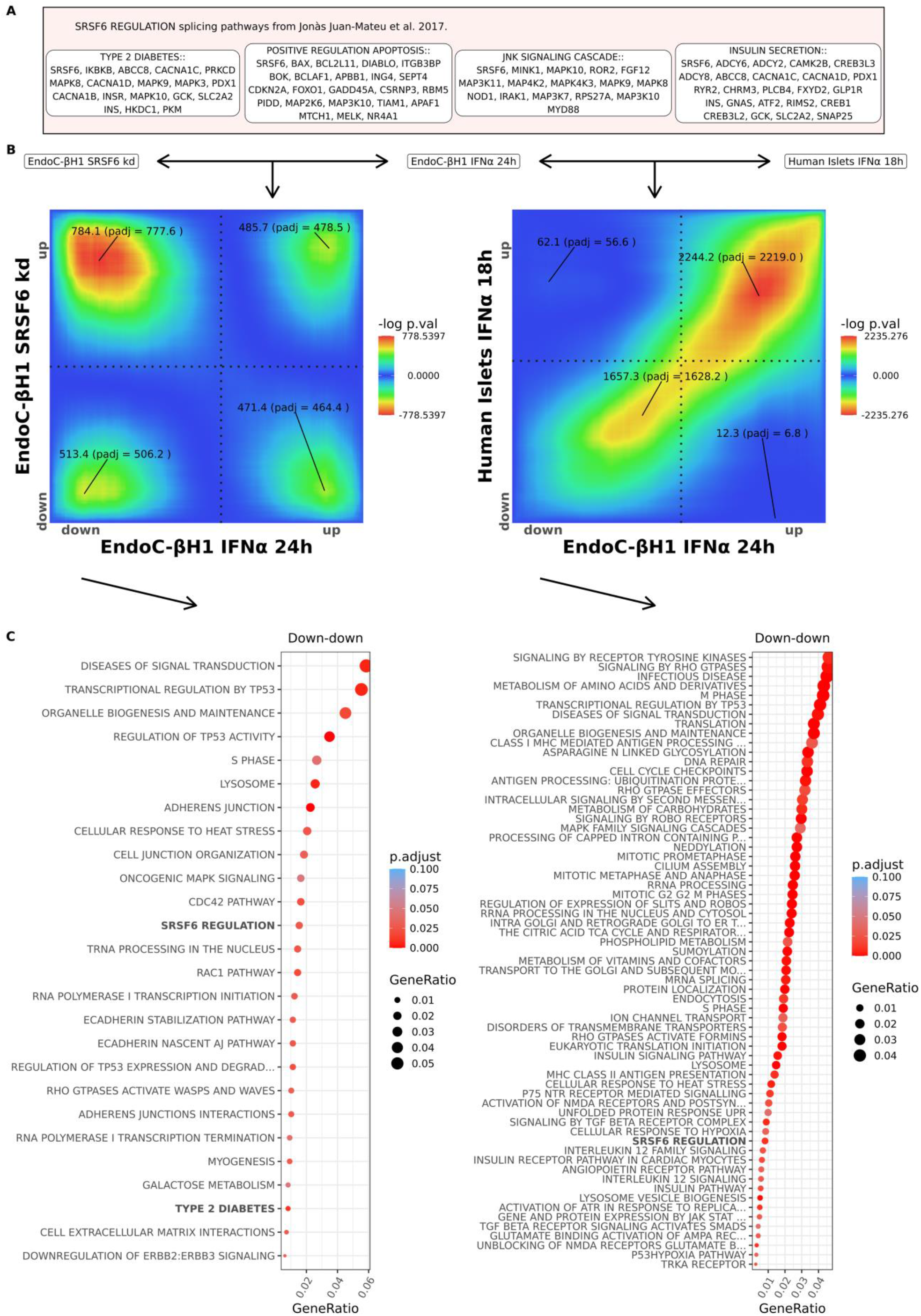
Transcript level analysis of SRSF6 regulated alternative splicing network. (A) Genes and pathways regulated by SRSF6 as identified in (Juan-Mateu, et al., 2018). (B) RedRibbon transcript level overlap maps comparing differential analyses of IFNα-treated and SRSF6-silenced EndoCβH1 cells (left), and IFNα-treated EndoC-βH1 cells and IFNα-treated human islets (right). (C) Molecular Signature Database and canonical pathways enriched in overlapping downregulated transcripts. The pathways known to be regulated by SRSF6 are highlighted in bold.

The transcripts upregulated in EndoC-βH1 cells by *SRSF6* knockdown and IFNα exposure exhibit significant enrichment in interferon signalling, lysosomes, and apoptosis (Fig. 5A). Overlapping upregulated transcripts between IFNα-exposed EndoC-βH1 cells and human islets showed additional enrichment of alternative splicing networks with a major role in beta cell signalling and apoptosis (Fig. 5B). Of note, 11 of the 16 enriched pathways in EndoC-βH1 cells (Fig. 5A) were also present in human islets (Fig. 5B, highlighted in bold). Hallmarks of beta cells in type 1 diabetes were enriched, including the triad of MHC class I, stress pathways and inflammation (toll-like receptors, interferon signalling).

**Fig. 5.**
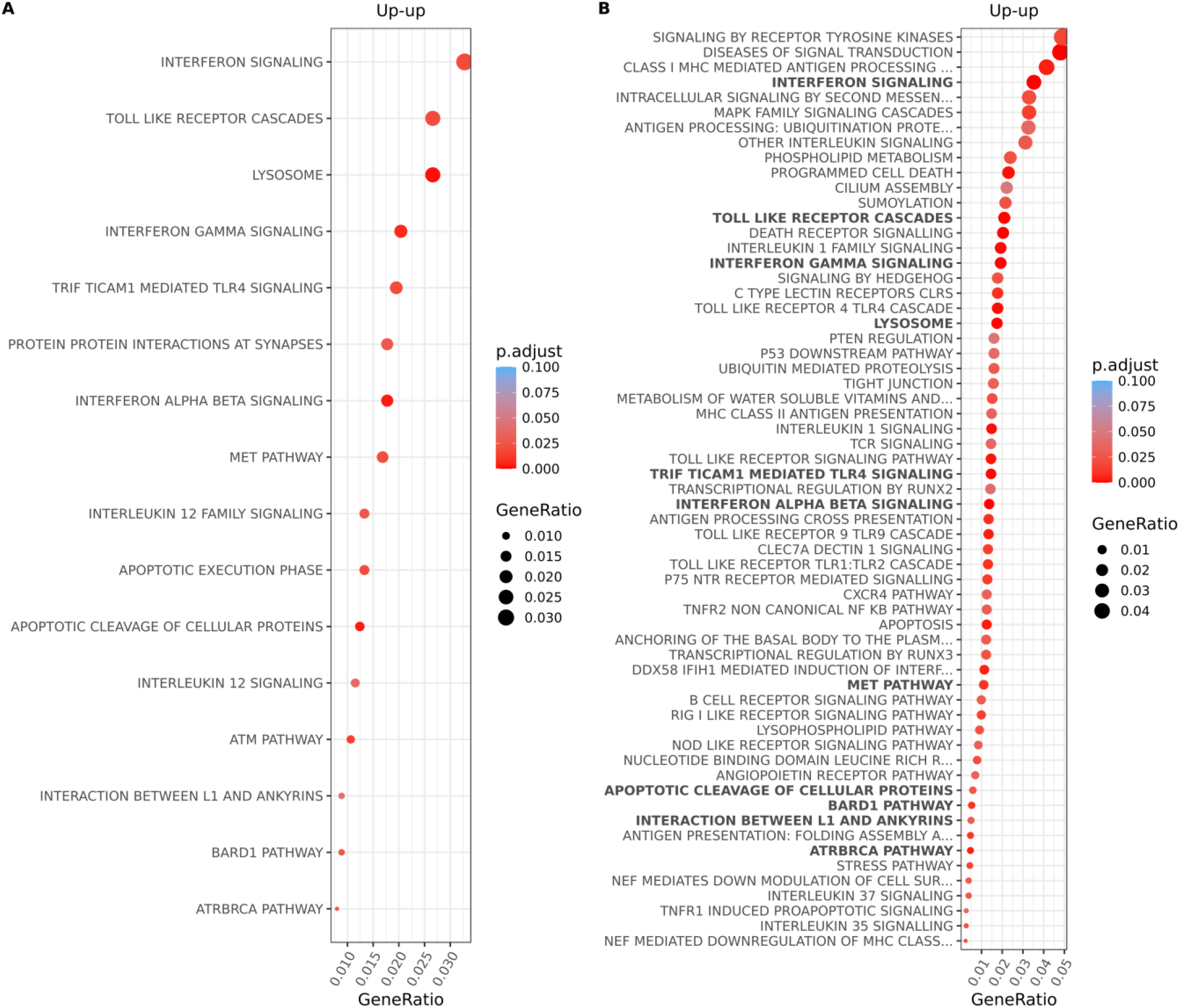
Molecular Signature Database and canonical pathways enriched in overlapping upregulated transcripts in beta cells following SRSF6 silencing and IFNα exposure. (A) Pathways enriched in upregulated transcripts in IFNα-treated and SRSF6-silenced EndoC-βH1 cells. (B) Pathways enriched in upregulated transcripts in IFNα-treated EndoC-βH1 cells and human islets. Pathways present in (A) are highlighted in bold in (B).

## 4 Discussion

The widespread use of omics technologies has exacerbated the need for new methods to analyse and compare diverse and ever larger datasets. Here we developed RedRibbon, a complete rewrite of the original RRHO package (Plaisier, et al., 2010), substantially increasing performance and accuracy, and introducing novel data structures and algorithms. In addition to the improved speed and accuracy of gene-level analyses, RedRibbon allows to leverage transcript-level quantification to detect overlapping signatures between two differential alternative splicing analyses. It features the capability to analyse lists one or two orders of magnitude longer without any loss of accuracy. The algorithms and data structures have been specifically tailored to be efficient (e.g., the bitset data structure allows to efficiently compute large set intersections using previously computed intersections). This new implementation goes beyond improving performance. First, gene or transcript overlap sets are provided for all four directions of regulation owing to the implementation of a two-sided test. The original R package did not report anticorrelated genes properly and the returned results were difficult to interpret (Cahill, et al., 2018). The updown and down-up quadrant enrichment lists are returned as the intersection of the set between the best coordinate and the quadrant corner.

Second, the accuracy of the localisation of the minimal *P*-value is improved over the grid method with an evolutionary algorithm. The grid method is pervasive among RRHO derivatives (Cahill, et al., 2018; Rosenblatt and Stein, 2014; Thind, et al., 2019) and other rank-based algorithms (Antosh, et al., 2013). For an analysis of 20,000 genes, the original R package grid used a default step size of 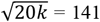, limiting the accuracy to this step size. The step parameter acts as a balance between speed and accuracy. A large step offers faster speed to the detriment of accuracy, and vice versa. Finding the right balance is difficult and there is no one-fits-all best value — the appreciation is left to the end user. As the error induced by the grid method is proportional to the step size and the complexity of the original R package algorithm is *O*(*n*(*n*⁄*step*)^2^) for transcript lists, a small step is computationally expensive while a large one gives inaccurate results leading to numerous false positives. Our evolutionary algorithm does not have this limitation and can accurately pinpoint the best *P*-value whatever the number of features analysed without impacting performance (Fig. 3). The evolutionary algorithm complexity is *O*(*i p n*) where *i* is the number of iterations (default value 200) and *p* is the population size (default value is 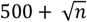 giving a default parameters complexity of 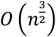. A downside is that it comes at the cost of nondeterminism in the minimal *P*-value finding algorithm. We mitigated this by initializing the algorithm population with evenly spaced coordinates on the diagonal of the map. In the experiments presently performed, we did not detect any minimal *P*-value worse than the ones detected by the grid method and the returned overlap sets were always close to identity in case of non-determinism.

Third, the computation of the overlap map is decoupled from the minimal *P*-value search. Hence, locating minimal *P*-value coordinates is independent of visualization map resolution. This helps to optimize memory usage, something that is particularly important in the analysis of very long lists. Our minimal *P*-value search algorithm only keeps in memory for the grid algorithm the best coordinates, and for the evolutionary algorithm the current population of coordinates, guaranteeing a very small memory footprint.

Eventually, a performant permutation scheme considering the correlation between genes is available to adjust the minimal *P*-value. This permutation scheme allows to correct the minimal *P*-value without having to rerun the whole differential analysis thousands of times while still considering the correlation between genes or transcripts. Doing so, the performance is greatly improved as shown in Fig. 3A. It makes it possible to run a permutation scheme over long lists and many conditions.

The package has been validated on synthetic datasets and previously reported RRHO results from (Marselli, et al., 2020) (Fig. 3B-E). The RedRibbon evolutionary algorithm detected synthetic dataset genes with a systematically better accuracy compared to the original algorithm. A similar difference in the number of detected genes was also present for real datasets related to type 2 diabetes suggesting similar accuracy improvements. The differences are particularly marked when the overlap signal is diffuse (e.g., Fig. 3D down-down quadrant) as the step size misses the minimum, the surrounding *P*-values being very close in a large area. The observed differences are propagated in the pathway analyses. Hence, pinpointing the minimal *P*-value with accuracy is an important and unique feature of RedRibbon.

The package has been further applied to and validated for previously reported alternative splicing results in EndoC-βH1 cells and human islets (Juan-Mateu, et al., 2018) (Fig. 4 and Fig. 5). These analyses were done at transcript level on lists comprising around 150,000 transcripts (see Supplemental Table S1), list lengths that are beyond the reach of the original R package. RedRibbon allowed to run these analyses with accuracy and permutation adjusted *P*-values in a matter of minutes. Our results suggest that SRSF6 splicing regulation transposes from EndoC-βH1 cells to human islets as the SRSF6 splicing pathways are enriched in both for downregulated transcripts. Moreover, most of the upregulated transcript pathways in EndoC-βH1 cells are recapitulated in human islets. The availability of human islets of Langerhans is limited, whereas EndoC-βH1 cells are readily available. The present analyses suggests that EndoC-βH1 cells recapitulate human islets alternative splicing patterns, making this cell line an appropriate model to study alternative splicing in human beta cells by deep sequencing (Hastoy, et al., 2018; Lawlor, et al., 2019; Scharfmann, et al., 2014; Tsonkova, et al., 2018).

The results obtained here show the importance of transcript level analyses in order to capture the effects of alternative splicing. We designed the SRSF6 regulatory pathway based on previous splicing analysis (Juan-Mateu, et al., 2018), and it is only detectable at transcript level. For other pathways, one of the challenges is that pathway databases are gene oriented. For those, transcript sets returned by RedRibbon can be converted to genes before pathway enrichment to compensate for the lack of transcript level pathway databases. Using this method, we obtained a large intersection between gene and transcript level pathway analyses and identified many new pathways for transcript level analyses. Obviously, the final enrichment is only as good as the pathway databases. Splicing network regulatory pathways may not be detected without specifically tailored databases, as done here for SRSF6. The creation of new transcript level pathway databases will enable refined alternative splicing analyses.

In conclusion, RedRibbon is a very useful novel tool to compare both gene level and transcript level differential analyses. Specifically, RedRibbon allows the detection of splicing networks. Using this tool, we documented large transcript level similarities between EndoC-βH1 cells and human islets. Worth of note, the method is robust even when the cell types are not matched, as is the case here for immortalized beta cells and bulk human islets that contain around 50% beta cells. RedRibbon will be a very useful addition to the bioinformatic toolsets for the analysis and comparison of diverse ever bigger datasets.

## Acknowledgements

We wish to thank Xiaoyan Yi for testing the *R* package.

